# Three-Dimensional Binocular Eye-Hand Coordination in Normal Vision and with Simulated Visual Impairment

**DOI:** 10.1101/232488

**Authors:** Guido Maiello, MiYoung Kwon, Peter J. Bex

## Abstract

Sensorimotor coupling in healthy humans is demonstrated by the higher accuracy of visually tracking intrinsically-rather than extrinsically-generated hand movements in the fronto-parallel plane. It is unknown whether this coupling also facilitates vergence eye movements for tracking objects in depth, or can overcome symmetric or asymmetric binocular visual impairments. Human observers were therefore asked to track with their gaze a target moving horizontally or in depth. The movement of the target was either directly controlled by the observer's hand or followed hand movements executed by the observer in a previous trial. Visual impairments were simulated by blurring stimuli independently in each eye. Accuracy was higher for self-generated movements in all conditions, demonstrating that motor signals are employed by the oculomotor system to improve the accuracy of vergence as well as horizontal eye movements. Asymmetric monocular blur affected horizontal tracking less than symmetric binocular blur, but impaired tracking in depth as much as binocular blur. There was a critical blur level up to which pursuit and vergence eye movements maintained tracking accuracy independent of blur level. Hand-eye coordination may therefore help compensate for functional deficits associated with eye disease and may be employed to augment visual impairment rehabilitation.

## Introduction

To track moving objects, the visual system executes *smooth pursuit* eye movements, which hold the high resolution fovea onto a tracked object by matching its speed [53]. Furthermore, because humans have binocular overlapping visual fields, to inspect objects at different distances or to track objects moving in depth the oculomotor system also executes *vergence* eye movements, which are unequal slow rotations of each eye that shift the binocular gaze point in depth [15]. Smooth pursuit and vergence eye movements have different tracking characteristics [81] and rely on separate neural substrates [25, 100], however, they are similarly encoded in the Frontal Eye Fields region of frontal cortex [24, 1].

To actively interact with the environment, humans not only move their eyes, but also perform a range of 3D hand movements which include reaching, grasping and manipulating objects. In order to take advantage of correlations that exist between spatial coordinates in visual and action space, the eye and hand motor systems have been found to be linked. For example, Steinbach and Held [86] showed that smooth pursuit eye movements to self-generated motion are more accurate than to externally-generated motion. This finding is consistent with the view that there is an exchange of information between the motor systems of the eyes and hands [76]. Furthermore, Chen et al. [13] provide convincing electrophysiological evidence that the oculomotor system may receive efference copy from the hand motor system. However, it is also possible that a common command signal controls coordinated hand and smooth pursuit eye movements [8].

In contrast to the findings regarding smooth pursuit eye movements, to-date no link has been found between hand movements and vergence eye movements in depth. Koken and Erkelens [46] in particular have shown that simultaneous hand tracking of a target moving sinusoidally in depth does not improve the latency of vergence eye movements executed to visually track the target. This finding is in contrast to previous results by the same authors which demonstrate that simultaneous hand tracking of an externally-generated target motion enhances smooth pursuit eye movements [45]. Thus the vergence system does not exhibit the same eye-hand coupling as the pursuit system when the eyes and hands are tracking externally generated motion. These findings suggest that the oculomotor system does not employ afferent hand motor signals to enhance vergence tracking. It remains unknown whether vergence eye movements may be enhanced during tracking of internally-generated motion in depth. If this were the case, it would provide evidence that either a common command signal controls coordinated hand and vergence eye movements, or that the oculomotor system receives efferent hand motor information when tracking self-generated motion in depth. Thus, the present study aims to investigate whether both pursuit and vergence eye movements may be enhanced during tracking of internally-generated motion.

Binocular vision and stereoscopic depth perception are fundamental in everyday eye-hand coordination [21, 67, 29]. Consequently, patients with stereovision deficits are known to have impairments in reaching and grasping hand movements (e.g. children with amblyopia [30, 88, 82] or adults with age related macular degeneration [98] or glaucoma [47]). One of the most basic and pervasive causes of visual impairment is refractive error (defocus blur), with an estimated 153 million visually impaired people worldwide as a result of uncorrected refractive errors [73]. Refractive error refers to the inability of the eye to focus light onto the retina. Myopia (nearsightedness) for example, is a common type of refractive error in which the eyes grow too large for the lens system to be able to focus far objects onto the fovea. Refractive error causes retinal images to be blurred and reduces the visibility of small objects and fine spatial detail. No official prevalence data are available, but for different age groups the distribution of myopic and hyperopic refractive errors is not negligible (see Figure 1 [84, 38, 42, 20]). Some visually-impaired individuals are defined by the type of refractive error: patients with anisometropic amblyopia tend to have significant hyperopic refractive error that is binocularly asymmetric, consequently untreated amblyopes have different amounts of blur in the two eyes [87].

**Fig. 1.**
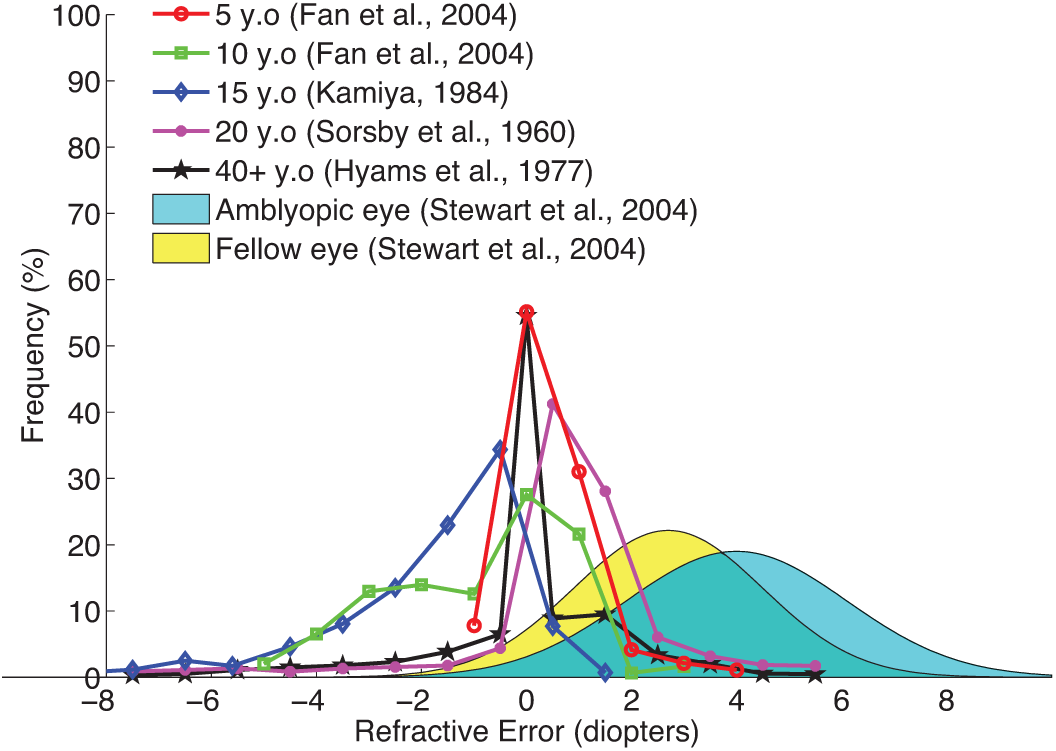
Refractive Error Distributions. Distributions of refractive errors for different age groups and for amblyopic children, replotted from a sample of the literature.

Symmetric and asymmetric visual blur may affect smooth pursuit and vergence eye movements in several different ways. Blur reduces both the contrast and the spatial frequency content of visual targets. Spering et al. [85] have found that reduced contrast impairs pursuit eye movements by affecting the correct estimation of target speed while reducing spatial frequency has no systematic effect on pursuit eye movements. Conversely, the precision of binocular alignment is known to depend on spatial frequency [78, 40, 54]. Thus, both smooth pursuit and vergence eye movements are likely to be impaired by symmetric blur in each eye. In conditions of asymmetric blur, unequal contrast and spatial frequency content in the two eyes may lead to suppression of the eye experiencing impoverished visual input [19, 49]. This interocular suppression is likely to have very different effects on the performance of smooth pursuit and vergence eye movements. During smooth pursuit eye movements, if the input from one eye were suppressed, pursuit performance might still successfully rely on the eye viewing the non-blurred target. Binocular disparity, a necessary component of vergence eye movements, is instead known to be impaired by interocular differences in spatial resolution[50]. Asymmetric blur is thus likely to impair vergence eye movements, because only the matching low spatial frequency information from the two eyes is available to maintain binocular alignment. Additionally, interocular suppression might mask the remaining matching binocular input altogether, making binocular alignment impossible.

However, it is still poorly understood how eye-hand coupling is affected by reduced target visibility. In conditions of visual uncertainty, the coupling between the eye and hand motor systems could break down. Conversely, von Noorden [66] showed that humans are able to make smooth pursuit eye movements to their own finger motion in the dark, therefore if the oculomotor system optimally combined visual, hand motor efferent and proprioceptive information, it is possible that eye-hand coupling could be unaffected, or even strengthened, when visual information is unreliable.

Here, we investigate the coupling between the eye and hand motor systems under conditions of simulated monocular and binocular blur. More specifically, we examine both left-right pursuit eye movements in the fronto-parallel plane as well as vergence eye movements in depth. Because eye hand coordination occurs in 3D, we ask how eye-hand coupling in depth is affected by blur, and whether monocular and binocular visual impairments have similar effects. We also discuss how our findings from image blur could relate to real-world blur associated with refractive error.

## Materials and Methods

### Participants

Five subjects, author GM and four naÏve observers, (3 male, mean ±sd age: 29±6) participated in the study. All subjects had normal or corrected to normal vision and normal stereo vision. All subjects reported being right handed and right-eye dominant. All procedures were approved by the Northeastern University Institutional Review Board and adhered to the tenets of the declaration of Helsinki. All subjects provided written informed consent.

### Apparatus

The experiment was programmed with the Psychophysics Toolbox Version 3 [10, 69] and Eyelink Toolbox [14] in Matlab (MathWorks). Stimuli were presented on an BenQ XL2720Z LCD monitor with a resolution of 1920 x 1080 pixels (display dot pitch 0.311 mm) at 120 Hz. The monitor was run from an NVidia Quadro K 420 graphics processing unit. Observers were seated in a dimly lit room, 57 cm in front of the monitor with their heads stabilized in a chin and forehead rest and wore active wired stereoscopic shutter-glasses (NVIDIA 3DVision) during all experiments to control dichoptic stimulus presentation. The cross talk of the dichoptic system was 1% measured with a Spectrascan 6500 photometer. Eye-tracking was performed using the EyeLink 1000 (SR Research) desktop mount eye-tracker running at 1000 Hz. The eye tracker was calibrated binocularly using the native five-point calibration routine at the start of each experimental session. Finger tracking was performed using a Leap Motion Controller, a commercial low-cost hand motion tracker recently validated for research applications [102, 31], which exhibits high accuracy (below 0.2mm) but an inconsistent sampling frequency of ≈ 40*Hz*. The Leap Motion device also exhibits a relatively high end-to-end latency (85ms) [12], that is nevertheless within a range that does not impair human performance at a range of tasks [101, 18, 91]. Binocular gaze and finger position measurements were queried from the eye and finger tracking devices at the monitor refresh rate of 120 Hz. Missing finger position samples were recovered in real-time through second order Savitzky-Golay interpolation on the previous 19 samples. The Leap Motion Controller was placed 30 cm in front and to the right of the observer, so that the observer’s right hand could be tracked. The Leap Motion was calibrated, once for all observers, so that placing the tip of the index finger 20 cm above the sensor was mapped to the center of the monitor, and left-right, backwards and forwards finger movements were mapped onto equally sized movements from the screen center. Left and right eye on-screen position data recovered from the Eyelink were transformed into the 3D gaze position with respect to the screen center.

### Stimuli

The stimulus for target tracking was a Gabor patch (*σ* = 0.25 degrees, *ω* = 2 cycles/degree, 100% Michelson contrast) moving on top of a white bar (0.5 degree thick and 8 degree wide). Stimuli were embedded in 1/f pink noise background which has the same frequency content of natural images [48, 7] and helped reduce the visibility of stereoscopic crosstalk. Example stimuli are shown in Figure 2. From trial to trial we systematically varied the amount of blur in the stimulus to one or both eyes using fast Gaussian mipmap filtering. Blur level is specified as double the standard deviation of the Gaussian blur kernel in minutes of arc. In the pursuit sessions seven blur levels were employed: [0, 6.25, 12.5, 25, 50, 100, 200] arcmin. In the vergence sessions only the first 6 blur levels were employed, as in pilot testing the vergence task was found to be already near impossible with 100 arcmin of blur. Note that with increasing blur levels the contrast of the stimuli decreased. The blur levels employed can be equated to contrast levels of [100, 95, 78, 59, 48, 35, 20] % Michelson contrast.

**Fig. 2.**
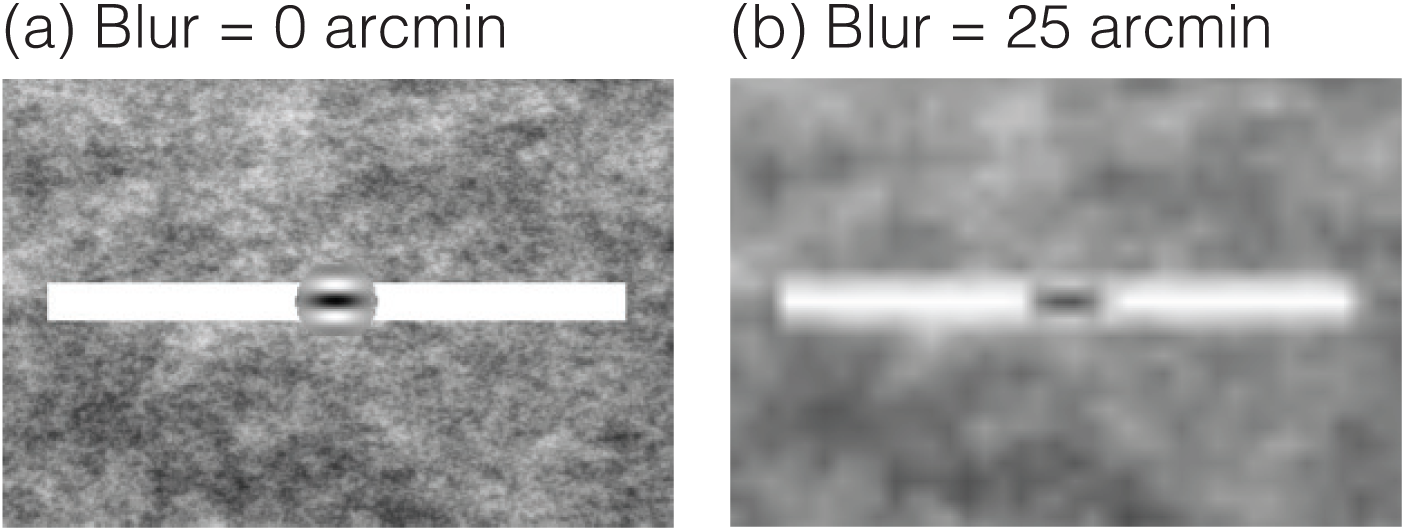
Example Stimuli. The stimulus for target tracking was a Gabor patch moving on top of a white bar. Background was 1/f pink noise. Left panel (a) is an example stimulus frame in the 0 arcmin blur condition. Right panel (b) is an example stimulus frame in the 25 arcmin blur condition.

In the binocular blur trials, the stimuli presented to both eyes were blurred. In the monocular blur trials, only the stimuli shown to the left eye were blurred, whereas a sharply focused stimulus was always shown to the right eye.

### Experimental Design

Observers participated in four experimental sessions on separate days. The four sessions are schematized in Figure 3. In sessions 1 and 2, which were horizontal pursuit eye movement sessions, observers were required to complete 5 trials for each of 7 blur levels for both monocular and binocular blur conditions. Thus in sessions 1 and 2 observers completed 70 trials per session. In sessions 3 and 4, which were vergence pursuit eye movement sessions, observers completed 5 trials for each of 6 blur levels for both monocular and binocular blur conditions, for a total of 60 trials per session. Within each session trial order was fully randomized.

**Fig. 3.**
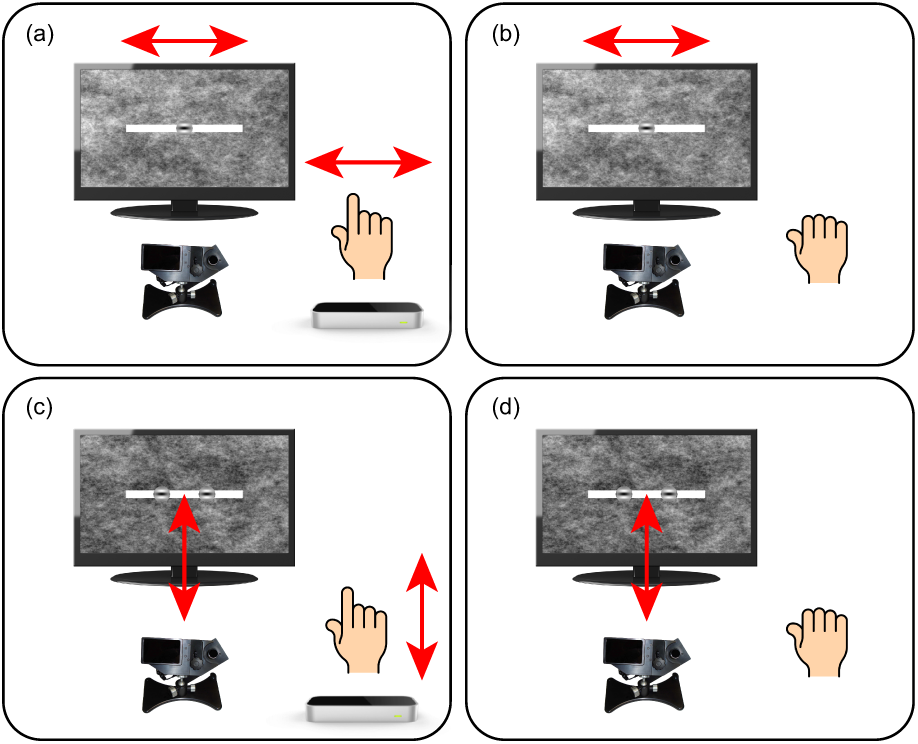
Schematics of the Four Experimental Sessions. In all four sessions observers’ binocular gaze point was monitored at 1000Hz with an Eyelink II eyetracker. (a) In session 1 observers smoothly moved their right index finger from right to left and back, directly above the Leap Motion Controller. The on-screen Gabor target moved with the observers’ finger. (b) In session 2 observers’ hands were motionless and the Gabor target moved right to left and back, replaying one of the finger movements executed in session 1. (c) In session 3 observers smoothly moved their right index finger backwards and forwards, directly above the Leap Motion Controller. The on-screen Gabor target moved in stereoscopic depth with the observers’ finger. (d) In session 4 observers’ hands were motionless and the Gabor target moved backwards and forwards, replaying one of the finger movements executed in session 3.

– Session 1: Horizontal Pursuit Finger Tracking (Figure 3a). At the beginning of each trial, observers placed their index finger above the Leap Motion Controller. When ready, observers pressed, with their left hand, the spacebar on the keyboard in front of them to begin the trial recording. Then, observers were required to smoothly move their index finger from right to left and back, directly above the Leap Motion Controller, which tracked the motion of the observer’s fingertip. The on-screen Gabor target moved with the observer’s finger, and the observer was required to track the Gabor target as accurately as possible with his/her gaze. The motion of the onscreen target was limited to within ±4 degrees from the center, constrained onto the stimulus bar. Once the observers had completed the movement, they signaled the end of trial recording by once again left-handedly pressing the space-bar on the keyboard in front of them. If less than 2 seconds passed between trial start and finish key-presses, the trial was deemed invalid and was repeated.
– Session 2: Horizontal Pursuit Replay Tracking (Figure 3b). Observers were required to hold their right hand motionless. When ready, observers pressed, with their left hand, the space-bar on the keyboard in front of them to begin each trial recording. Then, the Gabor target would move right to left and back, replaying one of the finger movements executed in session 1, matched to the viewing condition. The observer’s task was solely to track the Gabor target as accurately as possible with his/her gaze.
– Session 3: Vergence Pursuit Finger Tracking (Figure 3c). At the beginning of each trial, observers placed their index finger above the Leap Motion Controller. When ready, observers pressed, with their left hand, the spacebar on the keyboard in front of them to begin the trial recording. Then, observers were required to smoothly move their index finger backwards and forwards, directly above the Leap Motion Controller, which tracked the motion of the observer’s fingertip in depth. The on-screen Gabor target moved in depth with the observer’s finger, and the observer was required to track (by executing vergence eye movements) the Gabor target as accurately as possible with his/her gaze. The motion of the on-screen dot was limited from 0 to −8 degrees of crossed disparity (i.e. the dot could only move from the surface of the screen toward the observer). Once the observers had completed the movement, they signaled the end of trial recording by once again left-handedly pressing the space-bar on the keyboard in front of them. If less than 2 seconds passed between trial start and finish key-presses, the trial was deemed invalid and was repeated.
– Session 4: Vergence Pursuit Replay Tracking (Figure 3d). Observers were required to hold their right hand motionless. When ready, observers pressed, with their left hand, the space-bar on the keyboard in front of them to begin each trial recording. Each trial, the Gabor target moved backwards and forwards, replaying one of the finger movements in depth executed in session 3, matched to the viewing condition. The observer’s task was solely to track the Gabor target as accurately as possible with his/her gaze.

### Training

Prior to sessions 1 and 3, observers performed training to learn the eye-hand coordination task. Prior to session 1, observers performed 5 practice trials in which they moved their finger right to left and back, and were required to visually track the on-screen Gabor target. Prior to session 2, observers performed 5 practice trials in which they moved their finger backwards and forwards, and were required to visually track, via vergence eye movements, the on-screen Gabor target. Observers were informed that if less than 2 seconds passed between trial start and finish key-presses, the trial would be deemed invalid and have to be repeated. There was no upper time limit on the duration of a trial. In both sets of practice trials, visual stimuli were rendered with no blur to both eyes. Following the practice trials observers were informed that throughout the main experiment the visibility of the visual stimuli would vary.

### Statistical Analyses: A-Priori, Hypothesis-Driven Analyses

An initial aim of this study was to test whether both pursuit and vergence eye movements to self-generated hand movements are more accurate than eye movements to externally generated motion. We further test whether visual blur in one or both eyes disrupts visual tracking left right and in depth. Lastly, we test whether the link between the eye and hand motor systems could facilitate oculomotor target tracking in conditions of visual uncertainty.

To test our experimental hypotheses, analyses were designed a-priori by piloting our experimental design [58]. Pilot data showed that by measuring tracking accuracy as the correlation between gaze and target position we could reliably measure differences in accuracy as small as 1% for both pursuit and vergence eye movements. These pilot data also showed that the dependency of tracking accuracy as a function of blur was clearly not linear, but could be described by a horizontal asymptote at blur levels near zero and a steep fall-off at high blur levels. On linear-log axes, either a decreasing exponential function or its two-section piecewise linearization, a hinged-line function, described the pilot data equally well with the same number of free parameters. The hinged-line function was chosen over the decreasing exponential function because the fitted parameters lend themselves to a more immediate interpretation of the results, as we explain below. Thus, tracking data from our experiment was analyzed as follows.

Each trial, tracking accuracy was computed as the correlation between gaze position and target position either along the fronto-parallel plane (horizontal pursuit) or the sagittal plane (vergence pursuit). Figure 4 shows tracking accuracy as a function of blur for a representative observer in the binocular blur, horizontal pursuit finger tracking condition. These data were first transformed via Fisher's Z transformation to ensure variance stabilization [23]. Correlation data were then fit to a hinged-line linear-log function with equation:

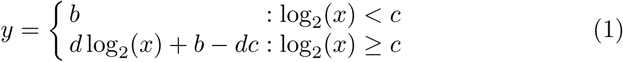

where *b* is an observer’s *baseline* level of tracking accuracy, *c* is the *critical blur* level at which tracking performance begins to deteriorate, and *d* is the rate of *decay* of tracking accuracy with increasing levels of blur beyond *c*.

**Fig. 4.**
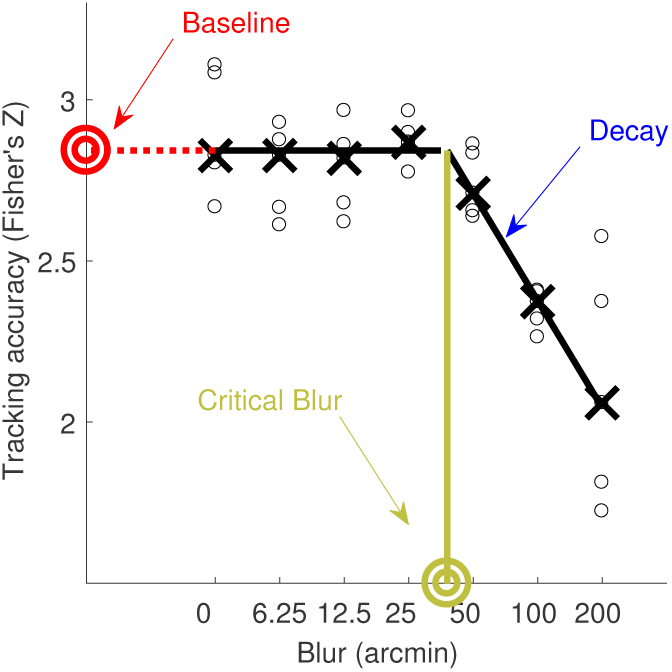
Example of Individual Tracking Data Fitted With A Hinged-Line Function. Tracking accuracy as a function of blur for a representative observer in the binocular blur, finger tracking, horizontal pursuit condition. Black circles are individual trial accuracy data. Black crosses are medians. Black lines are the fitted hinged-line linear-log function. The y-axis intercept is the *baseline* level of accuracy (red bullseye). Beyond the *critical blur* level (green bullseye), accuracy begins to falls below the baseline level of tracking. The slope of the linear fall-off (indicated by the blue arrow) is the rate of *decay* of tracking accuracy with increasing levels of blur.

If a link between eye and hand movements exists for both left-right and in-depth tracking, we would expect higher *baseline* accuracy in the finger tracking condition for both pursuit and vergence eye moments. If blur in one or both eyes impairs left-right and in-depth tracking differently, this would be reflected either by changes in the level of *critical blur* or by changes in the rate of *decay* of tracking accuracy. If the link between the eye and hand motor systems were to break down in conditions of visual uncertainty, we would expect either smaller *critical blur* or steeper *decay* in the finger tracking condition as compared the replay tracking condition. Conversely if eye-hand coupling were strengthened in conditions of visual uncertainty, we could expect larger *critical blur* levels or shallower *decay* rates during finger tracking.

To test these predictions, parameter estimates were analyzed using a 2 (Finger vs Replay Tracking) × 2 (Pursuit vs Vergence) × 2 (Binocular vs Monocular blur) within-subject Analysis of Variance (ANOVA). ANOVA normality assumptions were verified with Quantile-Quantile plots.

### Statistical Analyses: A-Posteriori, Exploratory Analyses

To tease apart the mechanisms underlying our primary findings, we proceeded to design the following exploratory analyses of our data. First, we processed the target trajectories of single trials from each experimental condition to obtain the onset of finger/target motion. We initially selected a rough maximum estimate of movement onset, which corresponded to the first position sample in which the target had moved in the direction of motion by at least 0.5 degrees for horizontal pursuit tracking and by at least 1 degree for vergence pursuit tracking. We then searched the movement traces up to this maximum estimate for the true movement onset using the method described by Schütz et al. [80, 13]. In short, velocity signals were calculated through digital differentiation, and regression lines with 50-ms length were fitted to these velocity trace. Regression lines with *R*^2^ < 0.7 or *slope* < 0.1 *degrees*/*s*^2^ were discarded. Of the remaining regression lines, the one with the highest *R*^2^ value was selected. The intercept of this line with the time axis was defined as the onset of the target motion. All target and eye movement traces from all experimental conditions were then temporally aligned so that time t=0 corresponded to the onset of the target motion. We then calculated duration and mean velocity of the finger movements, as well as the gaze-tracking error for every trial. Tracking error was computed as the gaze position minus the target position, along the frontoparallel plane for horizontal pursuit tracking and along the sagittal plane for vergence pursuit tracking.

To quantify the accuracy of open-loop tracking, i.e. tracking at movement onset, for each condition we estimated the onset of eye movements executed to track the target. We first low-pass filtered the eye position data below 10 Hz, then averaged the temporally-aligned eye position traces from the 5 repetitions for each condition. The onset of eye movements was then calculated using the same procedure employed to estimate target movement onset. The latency of horizontal and vergence tracking eye movements was thus defined as the difference between target and eye movement onset, with negative latencies indicating anticipatory eye movements. Latency data in the horizontal pursuit condition were analyzed using a 2 (Finger vs Replay Tracking) × 2 (Binocular vs Monocular blur) × 7 (Blur Level) within-subject ANOVA. Latency data in the vergence pursuit condition were analyzed using a 2 (Finger vs Replay Tracking) × 2 (Binocular vs Monocular blur) × 6 (Blur Level) within-subject ANOVA.

To quantify the accuracy of closed-loop tracking, i.e. tracking after movement onset, we instead measured the rate of saccadic eye movements during both horizontal pursuit and vergence tracking. Saccades were detected using a velocity threshold on the horizontal eye position traces of 25 degrees/s. Saccade rate was measured as the number of saccades occurring after the target had moved in the direction of motion by at least 0.5 degrees for horizontal pursuit tracking and by at least 1 degree for vergence pursuit tracking, divided by the duration of the trial. For horizontal pursuit tracking we included only catch-up saccades, i.e. saccades occurring in the same direction as the target motion. For vergence tracking we included all saccades occurring during the trial. Saccade rate data in the horizontal pursuit condition were analyzed using a 2 (Finger vs Replay Tracking) × 2 (Binocular vs Monocular blur) × 7 (Blur Level) within-subject ANOVA. Saccade rate data in the vergence pursuit condition were analyzed using a 2 (Finger vs Replay Tracking) × 2 (Binocular vs Monocular blur) × 6(Blur Level) within-subject ANOVA.

## Results

### Primary Findings from Hypothesis-Driven Analyses

Tracking accuracy varied lawfully as a function of blur level (Figure 5a,b); accuracy was constant *baseline* up to a *critical blur* level, after which accuracy fell off on linear-log axes with a linear rate of *decay.* This pattern held true for both horizontal pursuit eye movements (Figure 5a,b squares) and vergence pursuit eye movements (Figure 5a,b triangles).

**Fig. 5.**
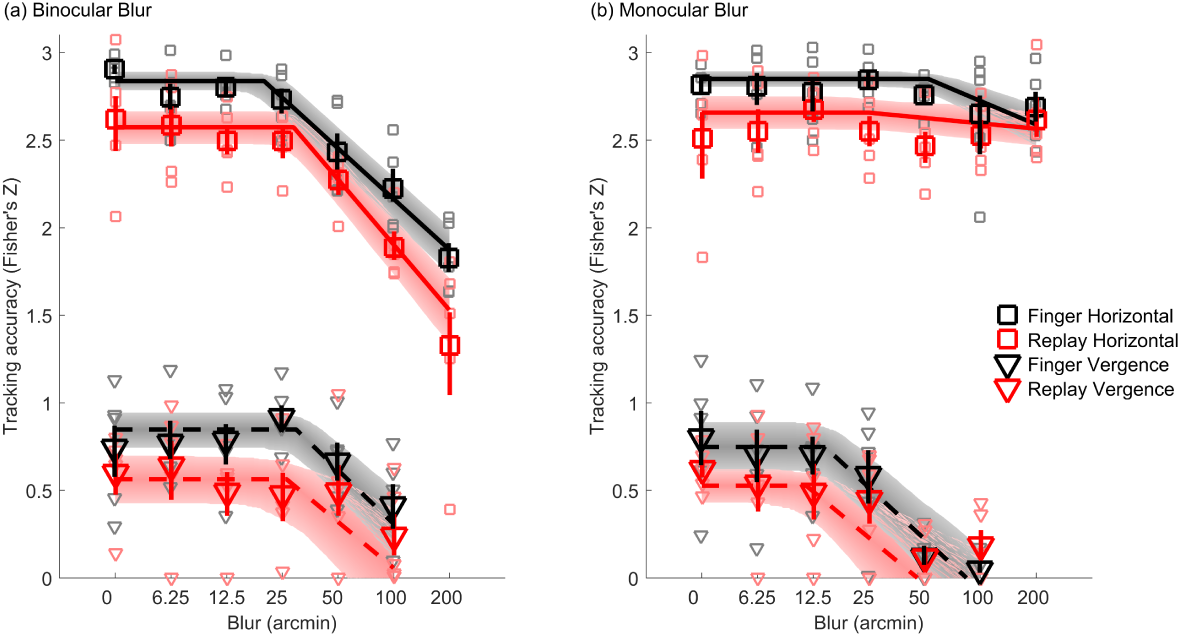
Tracking Accuracy. Tracking accuracy at each blur level for binocular (a) and monocular (b) blur conditions. Data from horizontal pursuit eye movements are shown as squares and vergence pursuit eye movements as triangles. Finger tracking data are shown in black and replay tracking are in red. Large symbols represent average tracking accuracy across subjects at each blur level. Error bars are 68% confidence intervals. Small symbols represent individual subject data. Solid and dashed lines are the average best fits of a hinged-line linear-log function for horizontal pursuit and vergence pursuit respectively. Shaded regions are 68% confidence regions of the fitted functions. Note that the y-axis is scaled following Fisher’s Z transformation, whereas the x-axis is log scaled.

#### 3D tracking of internally-generated motion is more accurate than tracking of externally generated motion

Tracking accuracy was noticeably higher when observers tracked a stimulus controlled by their own finger movements in real time (black curves, Figure 5a,b) than when observers tracked a replay of their own previously executed finger movements (red curves, Figure 5a,b). Specifically,Figure 6a shows that *baseline* accuracy was significantly greater in the finger (black) than the replay (red) tracking condition (Finger vs Replay main effect: *F*_1,4_ = 19.96, p = 0.011). Figure 6a also shows that horizontal pursuit (squares) tracking accuracy was three-fold greater (in Z-scaled space) than vergence pursuit (triangles) tracking accuracy (Horizontal vs Vergence main effect on *baseline* accuracy: *F*_1,4_ = 153.82, p = 0.00024). However, the difference between finger and replay conditions was similarly large across horizontal (Cohen’s d = 1.37) and in depth tracking (Cohen’s d = 0.94) conditions (Interaction effect on *baseline* accuracy between Finger vs Replay and Horizontal vs Vergence: *F*_1,4_ = 0.082, p = 0.79). This result further confirms the coupling between the oculomotor and hand motor systems and shows that the eye and hand motor systems are also linked for movements in 3D space. Sensibly, *baseline* accuracy did not vary as a function of monocular or binocular blur conditions, as these conditions are the same at baseline for the least blurred condition (Monocular vs Binocular blur main effect on *baseline* accuracy: *F*_1,4_ = 0.078, p = 0.79; all two and three way interactions: p > 0.16).

**Fig. 6.**
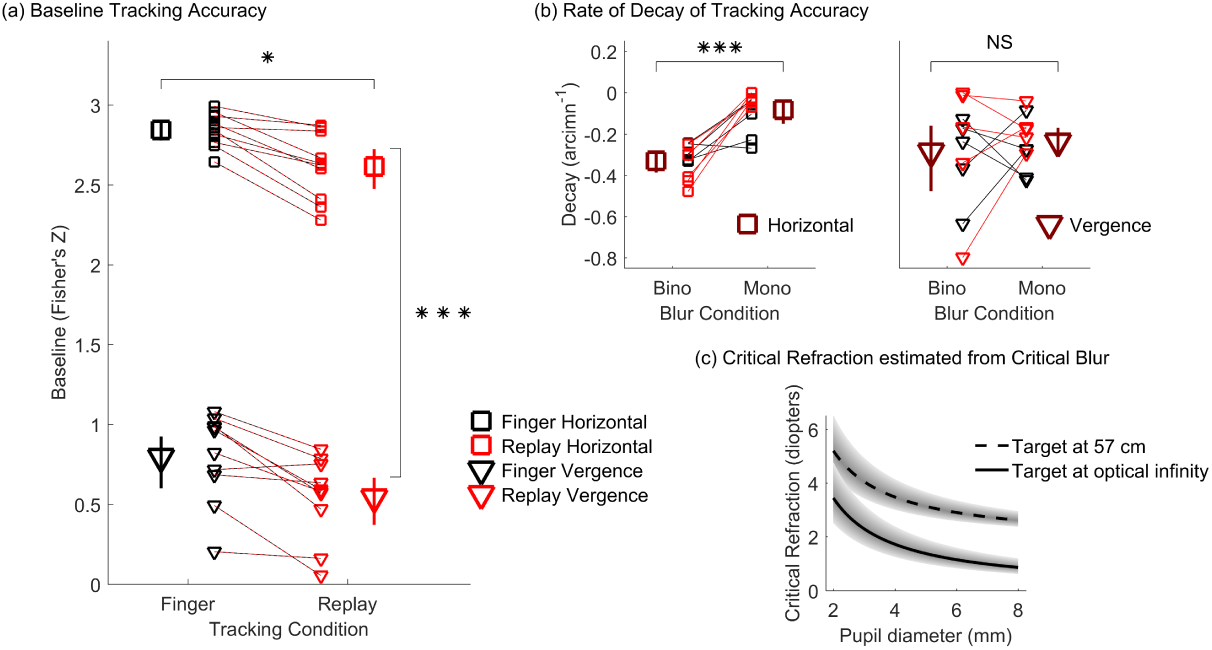
Primary Findings Based on the Fitted Parameters of the Hinged-Line Model. *Baseline* level (a), rate of *Decay* (b), and Critical Refraction (c) of tracking accuracy. In (a,b) black symbols represent finger tracking, red symbols represent replay tracking, squares are horizontal pursuit, triangles are vergence pursuit, small symbols represent individual subject data, and dashed red-black lines connect data points belonging to individual subjects from matching blur conditions. In (a) large black and red symbols are mean *Baseline* accuracy for the finger and replay conditions for both horizontal and vergence pursuit, averaged over monocular and binocular blur conditions. In (b) large brown symbols are the rate of *Decay* of tracking accuracy for the monocular and binocular blur conditions, averaged across both finger and replay tracking conditions, for both horizontal (left) and vergence (right) pursuit. All error bars are 95% confidence intervals. * p<0.05, *** p<0.001. (c) The estimated Critical Refraction (as computed from Eq. 2) at which oculomotor tracking performance may begin to decay is plotted as a function of pupil diameter for a target at 57 cm (dashed black curve) and for a target beyond optical infinity (continuous black curve). Shaded region encompasses the 95% confidence range of the estimate.

#### Eye-hand coupling is unaffected by blur

Tracking accuracy remained higher in the finger tracking condition compared to the replay tracking condition even when blur disrupted tracking performance. The rate of *decay* of tracking accuracy beyond the *critical blur* level did not vary between finger and replay tracking conditions (Finger vs Replay main effect: *F*_1,4_ = 0.53, p = 0.51). The
rate of *decay* of tracking accuracy was also similar for horizontal and vergence eye movements (Horizontal vs Vergence main effect: *F*_1,4_ = 1.23, p = 0.33).

#### Monocular blur hinders vergence tracking but not horizontal pursuit tracking

The rate of *decay* of tracking accuracy differed across monocular and binocular blur conditions (Monocular vs Binocular blur main effect: *F*_1,4_ = 43.04, p = 0.0028), and more specifically as a function of whether tracking was leftright or in depth (Interaction effect between Monocular vs Binocular blur and Horizontal vs Vergence: *F*_1,4_ = 9.87, p = 0.035). Figure 6b shows that when observers executed horizontal pursuit eye movements (left), the rate of *decay* was steep (i.e. more negative) if blur was rendered to both eyes and shallow if blur was rendered to one eye only, and this difference was statistically significant; t(4)= −10.23, p=0.00052, paired samples t-test, Cohen’s d = 5.97. When observers executed vergence eye movements instead, the rate of decay was similarly steep in both binocular and monocular blur conditions (as seen in the right panel of Figure 6b); t(4)= −0.99, p=0.38, paired samples t-test, Cohen’s d = 0.39. All other ANOVA two and three way interactions were not statistically significant (all p> 0.19).

#### Moderate amounts of blur impair tracking accuracy

The overall *critical blur* level at which tracking accuracy began to fall off was found to be 24 arcmin [18-32 arcmin, 95% confidence range]. This *critical blur* value was independent of the type of eye movement executed (Horizontal vs Vergence main effect: *F*_1,4_ = 1.41, p = 0.30), of whether observers were tracking their own finger movements or a replay of previously executed finger movements (Finger vs Replay main effect: *F*_1,4_ = 0.81, p = 0.42) and of whether blur was rendered monocularly or binocularly (Monocular vs Binocular blur main effect: *F*_i1,4_ = 1.031, p = 0.37; all two and three way interactions: p > 0.22).

The rendered blur employed in this study was meant to simulate the effects of uncorrected refractive error. A very simple approximate relationship exists between blur in angular units and dioptric level of defocus [83]:

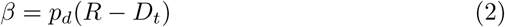

meaning that defocus in angular units *β* (radians) is equal to the pupil diameter *p_d_* (meters) multiplied by the difference between the refractive status of the eye *R* (diopters) and the distance to the target *D_t_* (diopters). Given that we have estimated the *critical blur* level at which tracking performance begins to deteriorate, we can employ Equation 2 to estimate at what level of refractive error oculomotor tracking performance begins to break down.

#### Common refractive errors may impair oculomotor tracking

Figure 6c shows the estimated Critical Refraction as a function of pupil diameter for a target at 57 cm (dashed line; within arm’s length) and for a target at optical infinity (continuous line; optical infinity in clinical practice is commonly defined as any distance farther than 6 meters). Pupil size is strongly dependent on light levels: pupils will constrict in high light and enlarge with low light levels. When visually tracking targets within arm’s length, only relatively large refractive errors (>5 diopters) will degrade oculomotor performance in outdoors, high light levels. In low light levels instead, 2.5 diopters of refractive error will already degrade oculomotor tracking of targets within arm’s length. When looking at optical infinity in high, outdoor light levels, up to 3 diopters of refractive error may be necessary to hinder oculomotor performance. For low, indoor light levels instead, 1 diopter of refractive error may already be sufficient to degrade oculomotor tracking performance for targets father than 6 meters away. Confronting these estimated values of critical refraction with the distribution of refractive errors found in the general population (Figure 1) highlights that a significant portion of children and adults may experience deficits in oculomotor tracking when refraction is not appropriately corrected.

### Complementary Findings from Exploratory Analyses

The findings from exploratory analyses help further characterize the differences in tracking performance between the finger and replay conditions under different experimental conditions. Figure 7 shows single trial tracking data from a subset of experimental conditions for one representative observer. In the horizontal pursuit tracking conditions (Figure 7a), individual trials lasted on average 3 seconds [2.5-3.6, 95% confidence range], and mean finger/target velocity was 5.4 deg/s [4.8-5.9, 95% confidence range]. Similarly, in the vergence pursuit tracking conditions (Figure 7b), individual trials lasted on average 2.9 seconds [2.4-3.1, 95% confidence range], and mean finger/target velocity was 5.8 deg/s [4.7-7.0, 95% CI]. It is obvious that the vergence data exhibits much more noise than the horizontal pursuit data, which is in line with the large difference in accuracy observed between the pursuit and vergence tracking conditions. Additionally, for both horizontal and vergence tracking, at all levels of monocular and binocular blur, the target (green curves) is more closely matched by the observer’s gaze position in the finger tracking condition (black) compared to the replay tracking condition (red curves).

**Fig. 7.**
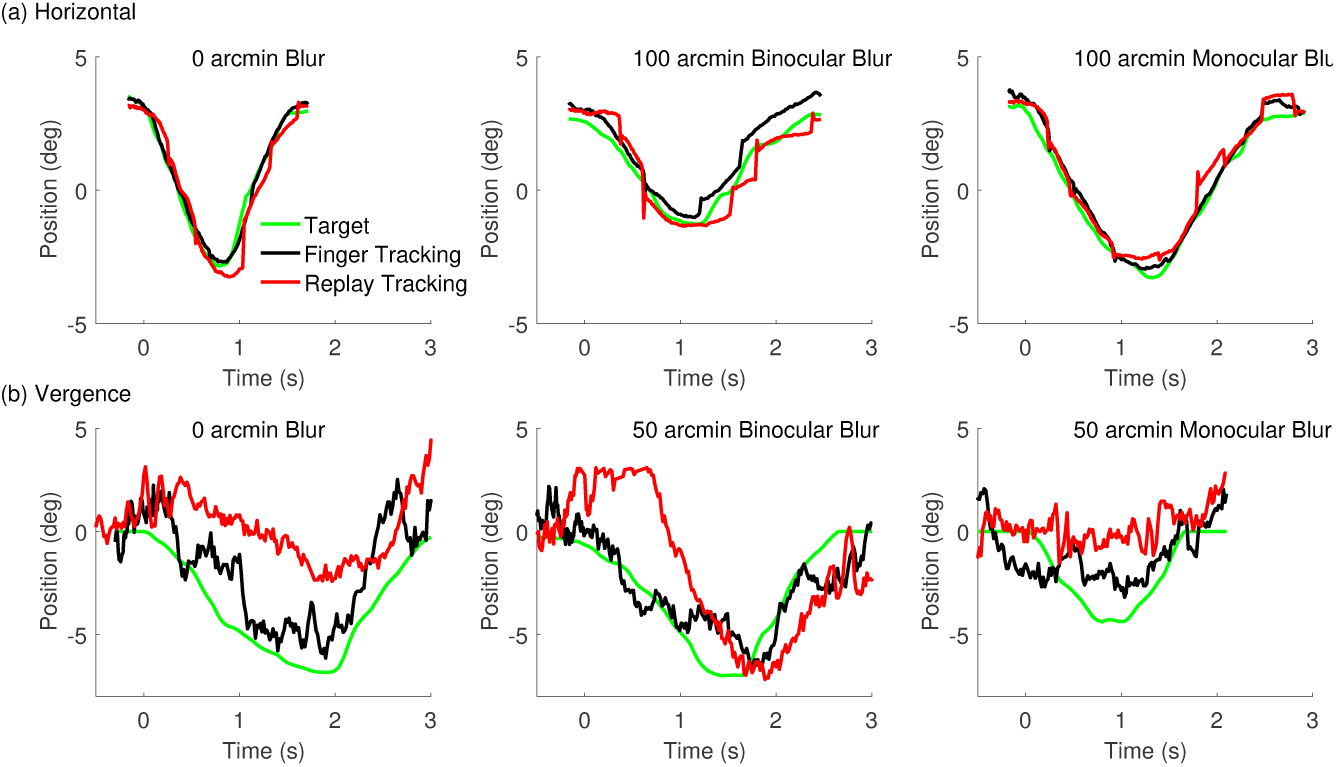
Raw tracking data from a representative observer. Target (green) and gaze position in the finger (black) and replay (red) tracking conditions are plotted as a function of time from target movement onset. (a) Horizontal pursuit tracking data from three trials. Left: a trial representative of the 0 arcmin blur condition. Center: a trial representative of the 100 arcmin binocular blur condition. Right: a trial representative of the 100 arcmin monocular blur condition. (b) Vergence pursuit tracking data from three trials. Left: a trial representative of the 0 arcmin blur condition. Center: a trial representative of the 50 arcmin binocular blur condition. Right: a trial representative of the 50 arcmin monocular blur condition.

#### Sensorimotor coupling aids the localization of the target

Figure 8 shows position error as a function of time from target movement onset for both horizontal pursuit (a) and vergence tracking (b), for the finger tracking condition in black and the replay tracking condition in red. The patterns of the error traces were qualitatively similar at all blur levels and in both the binocular and monocular blur conditions, with only the magnitude of the error varying in the different blur conditions. Thus, for simplicity we present the average error traces from all blur conditions. During horizontal pursuit tracking, (Figure 8a) observers anticipated the target movement onset (observe the positive deflection of the error traces at time zero), and this anticipatory deflection was more pronounced when observers were tracking their own finger (black curve) than when they were tracking a replay of their own previously executed finger movements (red curve). After movement onset, the tracking error remained markedly smaller in the finger tracking condition (black curve) than in the replay tracking conditions (red curve). During vergence pursuit tracking, (Figure 8b) observers did not visibly anticipate the target movement onset in either the finger or the replay tracking conditions. However, after target movement onset the tracking error was smaller during the finger tracking condition (black curve) than in the replay tracking conditions (red curve).

**Fig. 8.**
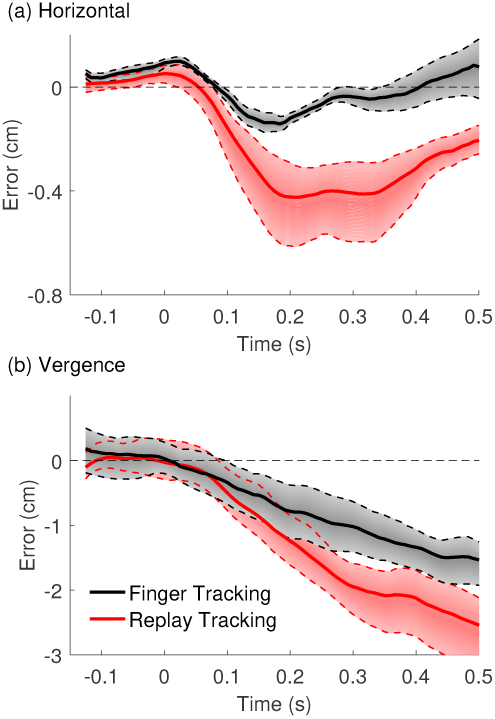
Time Course of Tracking Error. Tracking error (the difference between gaze and finger position) is plotted as a function of time from target movement onset for (a) horizontal pursuit and (b) vergence pursuit tracking. Positive error values signify that the eyes were leading the target, negative errors signify that the eyes were trailing the target. Black and red curves are the error in the finger and replay tracking conditions respectively, averaged across all monocular and binocular blur conditions. Shaded regions bounded by dotted lines are 68% confidence intervals of the mean error trace. Note that y-axes in (a) and (b) are scaled differently.

#### The eyes are nearly synchronous to the hand during horizontal pursuit tracking

To more closely investigate tracking performance at target movement onset, we measured the latency of gaze tracking in each experimental condition. Figure 9a shows how during horizontal pursuit tracking (black data), the eyes moved in almost perfect synchrony with the target during the finger tracking condition, whereas there was an average latency of ≈ 70ms in the replay tracking condition. ANOVA analysis confirmed that horizontal pursuit tracking exhibited a significantly shorter latency in the finger tracking condition compared to the replay tracking condition (Finger vs Replay main effect: *F*_1,4_ = 32.7466, p = 0.0046, Cohen’s d = 2.45). The latency of horizontal pursuit tracking did not vary as a function of blur level (Blur Level main effect: *F*_6,24_ = 0.70, p = 0.65), and was independent of whether blur was rendered monocularly or binocularly (Monocular vs Binocular blur main effect: *F*_1,4_ = 2.10, p =0.22). All ANOVA two and three way interactions were also not statistically significant (all p> 0.082).

**Fig. 9.**
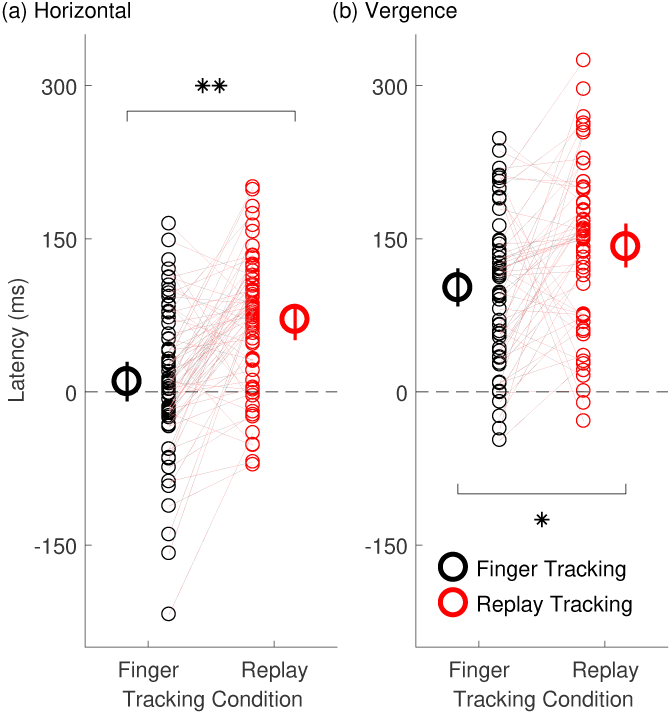
Tracking Latency. Latency of gaze tracking (i.e. the difference between target and eye movement onset in milliseconds) for (a) horizontal pursuit and (b) vergence pursuit tracking. Black circles represent finger tracking, red circles represent replay tracking, small symbols represent individual subject data, and dashed red-black lines connect data points belonging to individual subjects from matching blur conditions. Large symbols are mean estimates of tracking latency in the finger and replay tracking conditions, averaged across all monocular and binocular blur conditions. Error bars are 95% confidence intervals. Negative latencies indicate anticipatory eye movements. * p<0.05, ** p<0.01

#### Sensorimotor coupling facilitates the prediction of target movement onset also during vergence tracking

During vergence pursuit, as shown in Figure 9b, gaze tracking was always delayed with respect to the onset of the target motion. However, tracking latency was noticeably smaller during finger tracking (black data, on average ≈ 100ms) and larger in replay tracking (red data, on average ≈ 140ms). ANOVA analysis further confirmed statistically significant difference in vergence pursuit latency between finger and replay tracking conditions (Finger vs Replay main effect: *F*_1,4_ = 9.48, p = 0.034, Cohen’s d = 1.33). As with horizontal pursuit tracking, the latency of vergence pursuit tracking did not vary with blur level (Blur Level main effect: *F*_5,20_ = 0.51, p = 0.77) nor whether blur was rendered monocularly or binocularly (Monocular vs Binocular blur main effect: *F*_1,4_ = 0.021, p = 0.89). ANOVA analysis revealed no significant two or three way interaction (all p> 0.49).

#### Fewer catch-up saccades occur during tracking of self-generated target motion

To investigate errors arising from difficulties in perceiving the velocity of motion along the fronto-parallel or sagittal planes, we measured the rate of saccades occurring during each tracking trial. During horizontal pursuit tracking, we measured the rate of catch-up saccades that subjects executed to bring the two foveae back onto the target. Figure 10a shows that saccade rate was significantly smaller in the finger tracking condition (black) compared to the replay tracking condition (red; Finger vs Replay main effect: *F*_1,4_ = 37.11, p = 0.0037, Cohen’s d = 1.01). We found no significant main effect of Monocular vs Binocular blur conditions (*F*_1,4_ = 4.55, p = 0.10). There was however significant main effect of blur level (*F*_6,24_ = 5.77, p = 0.00078) and significant interactions effects between Finger vs Replay and Monocular vs Binocular conditions (*F*_1,4_ = 18.67, p = 0.012) and between Monocular vs Binocular conditions and blur Level (*F*_6,24_ = 7.86, p = 0.000093). These significant effects were all driven by the 200 arcmin, binocular blur condition, in which the average saccade rate fell significantly below the average saccade rate from all other experimental conditions (t(68) = −4.05, p = 0.00013, unpaired samples t-test; compare brown circle and star symbols in Figure 10a). There was no significant interaction between Finger vs Replay conditions and blur Level (*F*_6,24_ = 1.74, p = 0.15), nor a significant three way interaction (*F*_6,24_ = 1.06, p = 0.41).

**Fig. 10.**
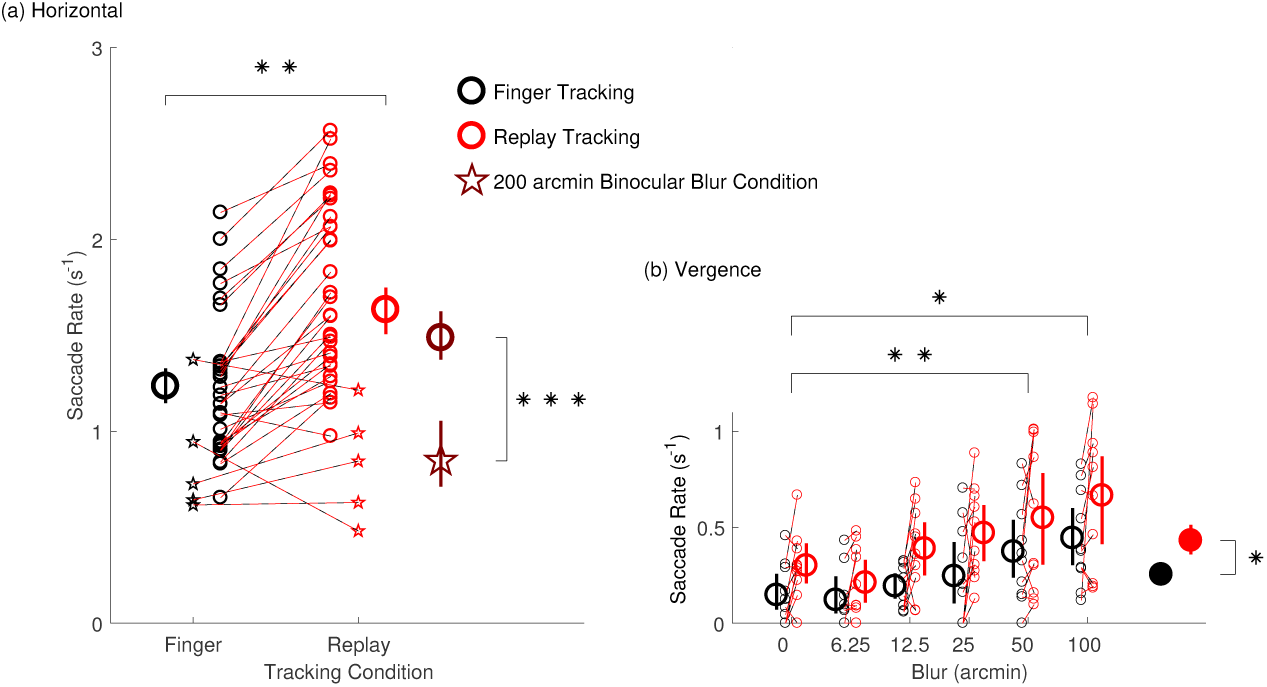
Saccade Rate. Number of saccades per second occurring during (a) horizontal pursuit and (b) vergence pursuit tracking. Black symbols represent finger tracking, red symbols represent replay tracking, small symbols represent individual subject data, and dashed red-black lines connect data points belonging to individual subjects from matching blur conditions. In (a) small star symbols are data from the 200 arcmin binocular blur condition, while small circles are data from all other conditions. Large black and red circles are mean latencies for the finger and replay tracking conditions respectively, averaged across all blur conditions. The brown star symbol represents the average saccade rate for the 200 arcmin binocular blur condition, whereas the brown circle represents the average saccade rate for all the other conditions. In (b) saccade rate during vergence tracking is plotted as a function of blur level. Large empty circles are mean saccade rate at each blur level, averaged across monocular and binocular blur conditions. Filled black and red circles are mean saccade rate during finger and replay tracking respectively, averaged across all blur conditions. All error bars are 95% confidence intervals. * p<0.05, ** p<0.01, *** p<0.001

#### Eye-hand coupling also increases the stability of vergence tracking

As expected, saccade rate in the vergence pursuit condition (Figure 10b) was lower than in the horizontal pursuit condition, as observers were required to track a stimulus which only moved in depth and not left-right. Nevertheless, Figure 10b shows how saccade rate varied as a function of blur level (Blur Level main effect: *F*_5,20_ = 17.16, p = 0.0000013), and specifically, saccade rate increased significantly from baseline when observers were presented with 50 arcmin (0 vs 50 arcmin blur: t(4) = −8.33, p = 0.0057, Bonferroni-corrected paired samples t-test, Cohen’s d = 1.51) and 100 arcmin of blur (0 vs 100 arcmin blur: t(4) = −6.00, p = 0.019, Cohen’s d = 2.76). Additionally, we found that saccade rate was statistically smaller during finger tracking than in the replay tracking (compare the black and red filled circles in Figure 10b) (Finger vs Replay main effect: *F*_1,4_ = 12.21, p = 0.025, Cohen’s d = 1.2832). However, no significant difference was found between monocular and binocular blur conditions (Monocular vs Binocular blur main effect: *F*_1,4_ = 2.11, p = 0.22) and no significant two or three way interaction effects were found (all p > 0.21).

## Discussion

We investigated the link between the oculomotor and hand motor control systems under binocularly asymmetric blur conditions, for both horizontal pursuit eye movements and vergence pursuit eye movements in depth. We replicated the classic results that smooth pursuit eye movements to self-generated horizontal motion are more accurate than to externally generated motion [86]. We extend these classic studies to show that the link between the eye and hand motor systems also exists for eye and hand movements in depth. Additionally, we show that moderate levels of simulated blur could disrupt oculomotor tracking performance, yet the link between the eye and hand motor system persists at all blur levels. Lastly, monocular and binocular blur have different effects on eye movements in the fronto-parallel (horizontal pursuit eye movements) and sagittal (vergence pursuit eye movements in depth) planes: Binocular blur similarly disrupts both horizontal pursuit and vergence pursuit eye movements. On the other hand, monocular blur strongly affects vergence eye movements while only mildly affecting horizontal pursuit eye movements.

To elucidate the potential mechanisms underlying our primary results we further characterized the patterns of tracking errors in our data using exploratory analyses. These analyses reveal that the linkage between the oculomotor and hand motor control systems helps predict the onset of target movement, and localize the target position during movement tracking. In other words, when observers tracked their own finger, tracking became more stable with anticipated eye movements (as shown by a reduction in both tracking latency and saccade rate), compared to when observers tracked an externally moving target. These patterns held true for both horizontal pursuit eye movements and vergence pursuit eye movements in depth.

### Mechanisms of Eye-Hand Coordination

#### Shared motor planning between the eyes and hands

It is possible that proprioceptive information about hand location may play a role in guiding eye movements [59, 60]. However, proprioception does not fully account for increased accuracy at tracking self motion, since in patients devoid of proprioception pursuit movements are still able to anticipate the start of target motion when the motion is self-generated [96]. Thus, it is more likely that the eye and hand systems share motor planning information that facilitates the initiation of tracking eye movements, and then hand and arm proprioception helps maintain oculomotor tracking accuracy throughout the execution of the eye movements [95, 97, 96, 51]. For example, a recent study [13] has shown that the lateralized readiness potential, which signals hand motor preparation, is associated with anticipatory smooth pursuit eye movements to self-generated finger motion. These findings provide convincing evidence that the hand motor system shares motor planning information with the oculomotor system: shared motor planning information could take the shape of a common command signal sent to both motor systems [8]. Alternatively, a coordination control system has been proposed to explain enhanced accuracy at tracking self motion, in which motor efference copy is employed to synchronize the oculomotor and hand motor systems [28]. This coordination control model is supported by the near-perfect synchrony of eye and hand movement onset in smooth pursuit studies, because if oculomotor and hand motor system received a common command signal, the differences (both biomechanical and neural) between these motor systems would lead to asynchronies in the hand and eye movement onsets [76]. In our study, horizontal pursuit eye movements were initiated almost exactly at the same time as the target motion in the finger tracking trials, which supports the notion that oculomotor tracking is synchronized with the target through hand motor efference copy. However, this did not hold true for vergence tracking eye movements, where the eyes always lagged behind the target motion onset even though tracking latency was reduced during finger tracking compared to replay tracking. The different latencies found for horizontal pursuit and vergenge tracking eye movements are consistent with known differences in the spatio-temporal characteristics of these different eye movements [81, 53, 105], and could reflect differences in the neural substrates underlying sensorimotor coupling for coordinated hand-eye movements in depth or along the frontal plane. The cerebellum, which is directly involved in the control of both pursuit [103, 92] and vergence [27, 26] eye movements, likely plays an important role in the synchronization of the oculomotor and arm motor systems [61, 62]. If the same synchronization signals from the cerebellum were to converge onto separate vergence and pursuit-specific neural loci [25, 100], these signals might differentially impact oculomotor tracking performance in depth and along the frontal plane.

#### Afferent signals from the hand reduce oculomotor tracking errors

The role of arm proprioceptive signal in maintaining oculomotor tracking accuracy is demonstrated by studies showing that, when visually tracking externally generated motion, manual tracking enhanced smooth pursuit eye movements (at least when the object motion is predictable) [45, 65]. Koken and Erkelens [46] have however shown that when observers executed vergence eye movements to track a stimulus moving predictably in depth under external control, concurrent hand tracking of the stimulus did not aid oculomotor tracking performance. These results suggest that the oculomotor system may employ limb proprioceptive (afferent) signals to plan eye movements in the fronto-parallel plane, but might be unable to do so for eye movements in depth. In the current study, we find that tracking accuracy is higher for both horizontal pursuit and vergence eye movements when observers track self-generated hand motion. More importantly, this finding is not only due to better predictions of movement onset, but also due to a better estimation of the target motion during tracking, as demonstrated by smaller tracking errors (Figure 8) and increased tracking stability (i.e. smaller saccade rate, Figure 10) in the finger tracking conditions for both pursuit and vergence tracking. Thus, our findings challenge previous findings [46] by suggesting that both efferent and afferent signals from the hand motor system may play a critical role in maintaining the accuracy of both horizontal pursuit and vergence tracking eye movements in depth.

#### The strength of sensorimotor coupling may remain invariant

If the oculomotor system were able to modulate the contributions of the visual input and the hand motor efferent/afferent signals when planning horizontal smooth pursuit and vergence tracking eye movements, we might expect the hand motor signals to be weighed more strongly in conditions of greater visual uncertainty. However, the rate of *decay* of tracking accuracy remained the same in the finger and replay tracking conditions. Taken together, the contribution of the hand motor signals to oculomotor planning appears to be invariant over the short times covered in the present study. We, however, cannot rule out the possibility that the contribution of motor efferent/afferent signals may change over longer periods of adaptation to visual impairment.

#### Only binocularly symmetric blur impairs fronto-parallel tracking

It should be noted that the rendered blur employed in our study reduced both the contrast and the spatial frequency content of the visual target. When blurred stimuli were presented to one eye only, it is possible that the interocular differences in contrast and spatial frequency content could trigger interocular suppression [19, 49]. Hence, we can speculate that with increasing blur level, the impoverished visual input to the blurred eye was likely to be suppressed. Previous studies have shown that reduced contrast impairs pursuit eye movements, whereas changes in spatial frequency have no consistent effect on fronto-parallel tracking performance [85]. Our data indeed show a marked reduction in accuracy for horizontal pursuit tracking at increasing binocular blur levels, consistent with what we expected from the reduced contrast of a target. On the other hand, horizontal pursuit tracking was not strongly affected by monocular blur, as oculomotor performance likely relied on the eye viewing the non-blurred target. In the case of horizontal pursuit tracking, suppression of the blurred eye might actually aid tracking performance by reducing the noise in the system. However, the strength of interocular suppression is known to fluctuate on short time scales [6]. Hence, our tracking data in the monocular blur, horizontal tracking condition (Figure 5b), which exhibited small fluctuations in accuracy (that were not well captured by the hinged-line model), might have been influenced by fluctuations in the strength of interocular suppression.

#### Both binocularly symmetric and asymmetric blur impair vergence tracking in depth

In vergence eye movements, the images projected onto the foveae of the two eyes need to be aligned and thus high spatial frequency content in the two eyes is required to be matched [78, 50, 40, 54]. The vergence system is also tightly linked to the accommodation system, which attempts to minimize visual blur at the fovea of both eyes [22, 40]. In particular, some pre-motor vergence neurons may attempt to combine disparity and blur information to drive vergence, and under conditions of either monocular or binocular blur these neurons will receive a mismatched drive from ‘blur’ and disparity [41]. Hence, in our study vergence tracking was always strongly impaired by high levels of blur because in both monocular and binocular blur conditions the matching low spatial frequency information from the two eyes was insufficient to drive accurate binocular alignment. Our statistical analyses show no significant differences in vergence tracking performance between monocular and binocular blur conditions. Nevertheless, Figure 5 shows an interesting pattern of results, in which a decrease in the performance of vergence eye movements appeared to be more pronounced at high levels of monocular blur as compared to at high levels of binocular blur. This pattern of data, which is the opposite to that observed for horizontal pursuit, is also consistent with suppression of the eye experiencing impoverished visual input. At high levels of monocular blur interocular suppression could mask the remaining matching binocular input altogether, making binocular alignment essentially impossible.

### Limitations and Future directions

We acknowledge some methodological limitations of our work which may be relevant to future research into sensorimotor coupling. In the current study, motion of the index finger determined visual target motion, yet the target was in a different position with respect to the finger. Additionally, the low-cost finger tracking device employed in the current study introduces a lag between the time when the finger actually moves and when the target motion moves on-screen. As such, these conditions may not faithfully represent the normal coordination between oculomotor and hand movement control systems. Nevertheless, our data and results are highly consistent with the previous findings reported by other groups[86, 28, 95, 45, 97, 96, 51, 76, 65, 13]. It should be also noted that our tracking latency data did not increase with increasing visual uncertainty (e.g., blur level in either horizontal or vergence pursuit tracking). However, this may be largely due to the predictability of the target movement, as previously suggested by Chen et al. [13]. Overall, our data suggest that eye-hand coordination need not be spatially and temporally aligned, and future work is needed to characterize the spatio-temporal limits of sensorimotor coupling.

Another technological limitation lies in the fact that shutter-glasses was used in the current study to render stereoscopic depth and to dichoptically vary the blur simulated in the two eyes. This could have impacted eye tracking data quality, since the eye-tracker viewed the observer’s eyes through the shutter-glasses. However, this method has been successfully used in our published gaze-contingent display[54] and saccadic adaptation [56] studies, both of which required highly accurate binocular eye movement recordings. In addition, simulating depth through shutter-glasses can potentially decouple vergence and accommodation [75, 37, 99], which might induce additional noise in the vergence-accommodation system. We, however, found that sensorimotor coupling survived this potential cue conflict. Nevertheless, the role of conflicting disparity and focus cues in virtual reality technology with respect to eye-hand coordination requires further investigation in relation with refractive error development [57], as we discuss further below.

It is also important to note that in the current study visual impairments were simulated by Gaussian blurring the visual input. It is possible that oculomotor performance may be affected by different kinds of blur (such as sinc blur which contains phase reversals typical of the modulation transfer function of an optical system with a circular aperture such as the human pupil [64]), and that observers may learn to adapt to the specific type of defocus blur arising from their own optics [3]. Future investigations should thus consider to examine hand-eye coordination and oculomotor control in the general population as well as monocular visual acuity for static optotypes. Furthermore, as our blur manipulation affect both the contrast and the spatial frequency content of the visual targets, in a future study it would be helpful to directly tease apart the contributions of contrast and spatial frequency to oculomotor control and eye-hand coordination in 3D.

A final note concerns the fact that all observers in our study were right eye dominant, and monocular blur was always presented to their non-dominant left eye. Interocular differences in contrast sensitivity and acuity due to eye dominance are measurable in normally sighted observers [49]. These differences are indeed known to affect stereoacuity [50], and to be associated with phoria, i. e., the amount of ocular deviation occurring when fixating a target with one occluded eye [68]. As binocular rivalry suppression is known to be skewed in favor of the dominant eye [34], a future study should consider the potential role of eye dominance.

### Clinical Significance of our Findings

#### Uncorrected refractive errors can significantly impair oculomotor performance

The *critical blur* level at which tracking accuracy began to decay was independent of all experimental manipulations in the present study. Furthermore, our additional analysis demonstrates that refractive errors between 1 and 5 diopters may be sufficient enough to produce this level of textitcritical blur, depending on the ambient light level and viewing distance. Consider this range of *critical refraction* with respect to the distribution of refractive errors found in the general population (Figure 1). A significant portion of the population has refractive errors that, in our study, correspond to a decrease in oculomotor tracking performance. Furthermore, individuals with amblyopia or those with binocularly asymmetric visual impairment (such as cataracts), not only have an average acuity loss beyond this level of *critical blur*, but even the average difference in refraction between the amblyopic and fellow eye (1.3 diopters) might be sufficient to impair oculomotor performance. Our findings are thus in line with multiple studies showing that eye-hand coordination skills are significantly impaired in children with amblyopia as compared to normal cohorts [30, 88, 82].

#### Binocular eye movements and visual processing may synergistically degrade in amblyopia

More importantly, our findings may help us better understand the aetiology of amblyopia. It has been thought that this visual dysfunction develops when binocular vision becomes decorrelated owing to either ocular misalignment or interocular difference in refractive error, in which the brain learns to suppress visual signals from the weak eye to avoid diplopia (double vision) [104]. More specifically, uniocular blur in early life has been shown to induce amblyopia in macaque monkeys by impairing the spatial resolution and contrast sensitivity of the affected eye[43, 35]. Importantly, uniocular blur also reduces the degree of binocular interaction throughout the visual pathways, a necessary component of binocular oculomotor control[63]. However, the development of oculomotor and perceptual deficits in amblyopia might be interdependent. If the visual input from one eye is not correlated with the other eye, our findings show that oculomotor control might begin to degrade. Vergence eye movements are likely to become less accurate [78, 40, 54, 55], but also the mechanisms that calibrate the accuracy of binocular saccades might begin to fail [2, 79, 56]. If this occurs during early life, when vision and oculomotor control are in development [4, 5, 74, 33, 32], oculomotor and perceptual deficits might exacerbate each other. Binocularly imbalanced visual input may lead to poor binocular oculomotor control. Impaired oculomotor control in terms of imprecise binocular alignment, inaccurate binocular saccades, and incorrect eye posture (ocular deviation) will further decorrelate the visual input to the two eyes. This increased decorrelation between the visual input to the two eyes may lead to increased suppression of the weak eye. Increased suppression may in turn exacerbate oculomotor deficits, and so on. Thus not only may the perceptual deficits occurring in amblyopia lead to oculomotor deficits: perceptual and oculomotor deficits together might form a feedback loop of visuomotor impairment.

#### Eye-hand coordination for visual rehabilitation

This work also has potential implications for visual rehabilitation strategies in which binocular vision and eye movements are impaired, such as amblyopia, strabismus and convergence insufficiency. Classical patching therapy might be counterproductive for the development of coordinated vergence eye movements because it forces the two eyes to work independently and does not favor the correct development of binocular oculomotor control. Dichoptic therapies that attempt to balance the input to the two eyes and favor conjugate eye movements might be better suited to ensure binocular cooperation [36, 93, 44, 52, 89, 90, 56, 9]. Here, we have shown that the link between the oculomotor and hand motor system is unaffected by simulated visual impairment. Thus, eye-hand coordination tasks [94] might be able to provide a boost to visual rehabilitation strategies by enhancing oculomotor control both along the fronto-parallel plane and particularly in depth. This tantalizing idea is further supported by the notion that visual processing in general, not solely oculomotor performance, is enhanced near the hand (which suggests some cross-modality overlap of spatial representations). For example, several studies have demonstrated that visual stimuli placed near the hands are more readily detected [16, 72, 39, 70], importantly even in visually impaired neurological patients [17, 77, 11]. Perry et al. [71] have shown that the orientation selectivity of V2 neurons is enhanced for stimuli near the hand via the sharpening of orientation tuning. The effects of hand proximity have not yet been thoroughly investigated for other perceptual dimensions, such as visual processing of motion and depth, which are highly relevant in binocular visual deficits. Future studies should therefore focus on determining whether hand proximity may favor binocular cooperation. If this were the case, then eye-hand coordination task could be employed to synergistically enhance both binocular oculomotor performance and binocular visual perception, thus significantly augmenting current visual rehabilitation strategies.

## Author contributions statement

G.M., M.K. and P.J.B. conceived and designed the study. G.M. programmed the experiments, collected and analyzed the data. All authors wrote the manuscript. The authors declare that they have no conflict of interest. All data files and analysis scripts are available from the Zenodo database (10.5281/zenodo.1100971).

## Acknowledgements

The authors thank Dr. Jing Chen for help with setting up the pursuit onset detection algorithm originally developed by Dr. Alexander Schtz, as well as two anonymous reviewers. This research was supported by National Institutes of Health grant R01EY021553.

